# Phylogenetics analysis of TP53 gene in humans and its use in biosensors for breast cancer diagnosis

**DOI:** 10.1101/2020.07.25.221440

**Authors:** Sara da Silva Nascimento, Pierre Teodósio Félix

## Abstract

Biosensors are small devices that use biological reactions to detect target analytes. Such devices combine a biological component with a physical transducer, which converts bio-recognition processes into measurable signals. Its use brings a number of advantages, as they are highly sensitive and selective, relatively easy in terms of development, as well as accessible and ready to use. Biosensors can be of direct detection, using a non-catalytic ligand, such as cell receptors and antibodies, or indirect detection, in which there is the use of fluorescently marked antibodies or catalytic elements, such as enzymes. They also appear as bio-affinity devices, depending only on the selective binding of the target analyte to the ligative attached to the surface (e.g., oligonucleotide probe). The objectives were to evaluate the levels of genetic diversity existing in fragments of the TP53 gene deposited in molecular databases and to study its viability as a biosensor in the detection of breast cancer. The methodology used was to recover and analyze 301 sequences of a fragment of the TP53 gene of humans from GENBANK, which, after being aligned with the MEGA software version 6.06, were tested for the phylogenetic signal using TREE-PUZZLE 5.2. Trees of maximum likelihood were generated through PAUP version 4.0b10 and the consistency of the branches was verified with the bootstrap test with 1000 pseudo-replications. After aligning, 783 of the 791 sites remained conserved. The maximum likelihood had a slight manifestation since the gamma distribution used 05 categories + G for the evolutionary rates between sites with (0.90 0.96, 1.00, 1.04 and 1.10 substitutions per site). To estimate ML values, a tree topology was automatically computed with a maximum Log of −1058,195 for this calculation. All positions containing missing gaps or data were deleted, leaving a total of 755 sites in the final dataset. The evolutionary history was represented by consensus trees generated by 500 replications, which according to neighbor-join and BioNJ algorithms set up a matrix with minimal distances between haplotypes, corroborating the high degree of conservation for the TP53 gene. GENE TP53 seems to be a strong candidate in the construction of Biosensors for breast cancer diagnosis in human populations.

## 1. Introduction

Biosensors are small devices that use biological reactions to detect target analytes (WANG, 2000). Such devices combine a biological component, which interacts with a target substrate, to a physical transducer, which converts bio-recognition processes into measurable signals (WANG, 2000; PATHAK *et al,* 2007). Its use brings a number of advantages, as they are highly sensitive and selective, relatively easy in terms of development, as well as accessible and ready to use. However, there are certain limitations, such as electrochemically active interferences in the sample, little long-term stability, and electron transfer problems (MEHRVAR; ABDI, 2004; SONG *et al,* 2006).

Biosensors can be direct detection (direct detection sensor or non-reticulated system), in which biological interaction is measured directly, using a non-catalytic ligand, such as cell receptors and antibodies, or indirect detection (marked sensor or reticulated system), in which there is the use of fluorescently marked antibodies or catalytic elements, such as enzymes. The crosslinked system has greater stability and is simpler to use, but the non-reticulated system has better sensitivity, shorter operating time and lower costs (MEHRVAR; ABDI, 2004; PATHAK *et al,* 2007; LIU *et al,* 2009). There are two types of biosensors, depending on the nature of the recognition event. Bio affinity devices, which depend on the selective binding of the target analyte to the ligand attached to the surface (e.g., antibody or oligonucleotide probe) and bioanalytical devices, in which an immobilized enzyme is used for target substrate recognition (WANG, 2000). Based on this information, the objective of this work was to present a review of bibliography describing the structure, functioning and applicability of biosensors in various technological areas.

## 3. Objective

### 3.1 General

To evaluate the levels of genetic diversity existing in fragments of the TP53 gene deposited in molecular databases.

### 3.2 Specifics

Evaluate the levels of polymorphism in the gene encoding the TP53 protein and develop methodologies that allow the investigation of patterns of genetic variability for this gene.

## 4. Methodology

### 4.1. Dataset

Initially, 301 sequences of a fragment of the human TP53 gene recovered from GENBANK (https://www.ncbi.nlm.nih.gov/popset/430765060) and participated in a PopSet made available by Hao, X.D and collaborators in 2013 (PopSet: 430765060) were recovered and analyzed.

### 4.2. Analyses

After alignment with the mega software version 4.0 (KUMAR *et al.*, 2007), the phylogenetic signal will be tested using TREE-PUZZLE 5.2 (SHIMIDT, 2002). Trees of maximum likelihood will be generated through PAUP version 4.0b10 (SWOFFORD, 2002) and to evaluate the consistency of the branches, the bootstrap test (FELSENSTEIN, 1985) with 1000 pseudo-replications will be used. For the visualization of variable sites, logos will be generated through the Weblogo3 program (CROOKS, 2004). The analysis of the number of populations will be performed with the Structure 2.3 program (PRITCHARD, 2000) and two different methods are tested: a posteriori probability and ad hoc (k). The *“a posteriori”* probability will be calculated using an ancestry model with mixed alleles for 20,000 interactions in the burn-in period, followed by 200,000 Monte Carlo interactions via Markov Chain, increasing only the K value (number of populations), which will be from 1 to 10 according to Pritchard’s methodology (2000).

The Evanno method (2005) will be used to determine the most appropriate number of populations for the dataset, using an ad hoc amount based on the second-order rate of the likelihood function between the successive values of K. Posteriori and k probability tests will initially be applied to the dataset in isolation. For the analysis of genetic variability, a project will be created with the Arlequin Software 3.1 (EXCOFFIER *et al.,* 2005). which aims to measure molecular diversity using standard estimators such as Theta (Hom, S, k, Pi), Tajima Neutrality test, paired and individual F_ST_ values, in addition to temporal divergence and demographic expansion indices (mismatch and Tau values) by molecular variance analysis (AMOVA) (EXCOFFIER, 1992). In this method, the distance matrix between all haplotype pairs will be used in a hierarchical variance analysis scheme producing estimates of variance components analogous to Wright’s F statistics involving nonlinear transformations of the original information in estimates of genetic diversity. Mantel’s Z statistic will be used to represent the divergence between possible microhabitats using the MULTIVAR (Mantel for Windows) program (MANTEL, 1967).

## 5. Results

After being aligned, 783 of the 791 sites remained conserved. The maximum likelihood had a discrete manifestation for the gamma distribution with 05 categories + G for the evolutionary rates between sites with 0.90 0.96, 1.00, 1.04 and 1.10 substitutions per site. Nucleotide frequencies were A = 24.37%, T/U = 22.12%, C = 23.58% and G = 29.93%. For ML values, a tree topology was automatically computed with a maximum Log of −1058,195 for this calculation (Figures 1a and 1b). All positions containing missing gaps or data were deleted, leaving a total of 755 sites in the final dataset.

**Figure 1a.**
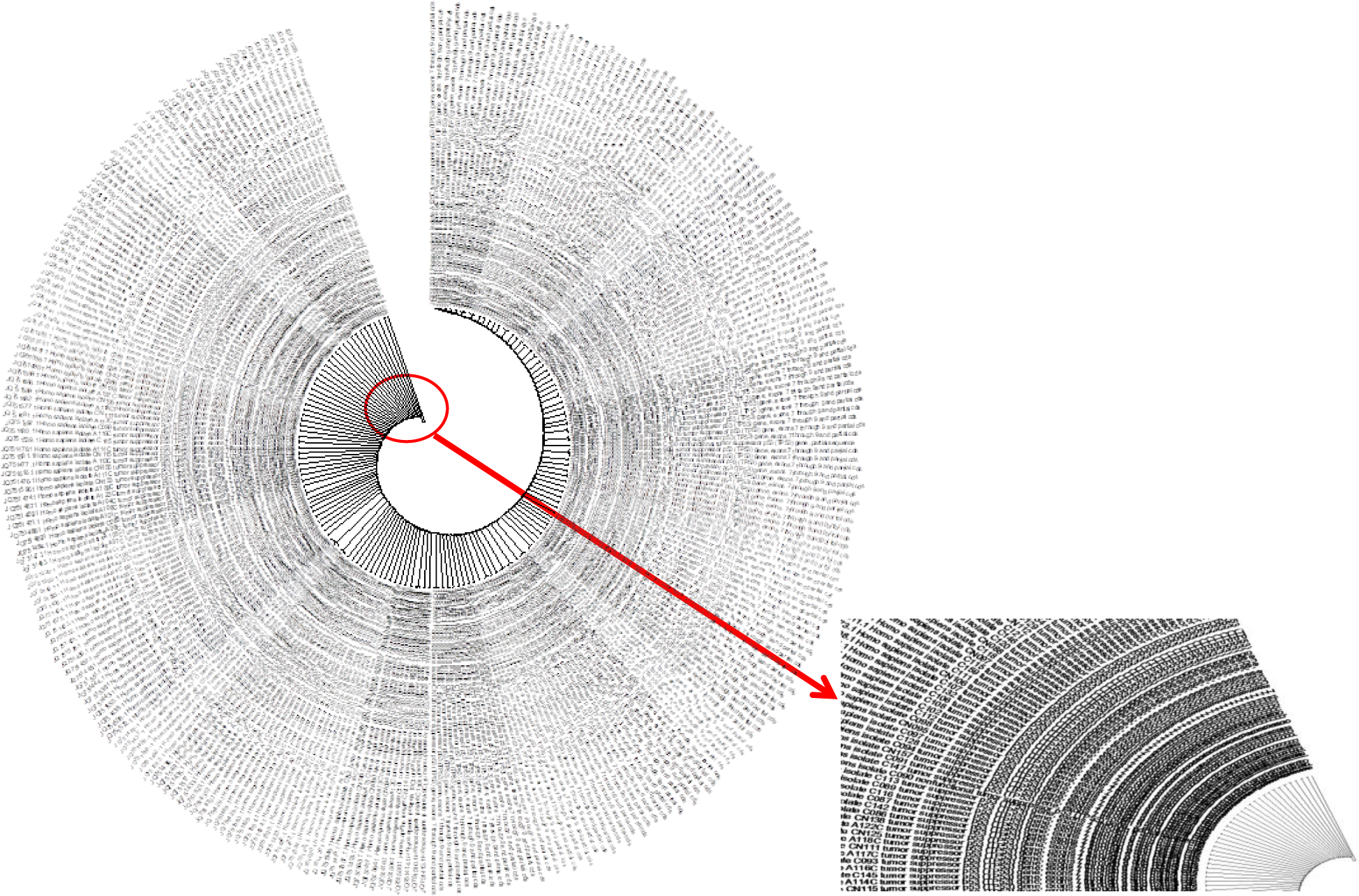
The analysis involved 301 nucleotide sequences. The codon positions included were 1st + 2nd + 3rd + Non-coding. All positions containing gaps and missing data were eliminated. There was a total of 755 positions in the final dataset. Evolutionary analyses were performed in MEGA6. **Figure 1b.** Cut showing the details of the haplotypes in the ML tree.

The evolutionary history was represented by consensus trees generated with 1,000 replications, which according to the algorithms of Neighbor-Join and BioNJ, set up a matrix of distance between the haplotypes that corroborated the high degree of conservation for the gene. For molecular variance tests, the 301 sequences were divided into 07 groups (04b, 05c, c85, 98c-1, a9cl, cn160 and a125c) that did not present levels of molecular diversity (0.05) (figure 2a, 2b), as well as in the Ewens-Watterson, Chakraborty, Tajima D and Fu Fs tests (table 1). In the FST tests, the only important variations were found within groups c85 and 04b with 0.73 and 0.39 respectively (figure 3, figure 4).

**Figure 2a.**
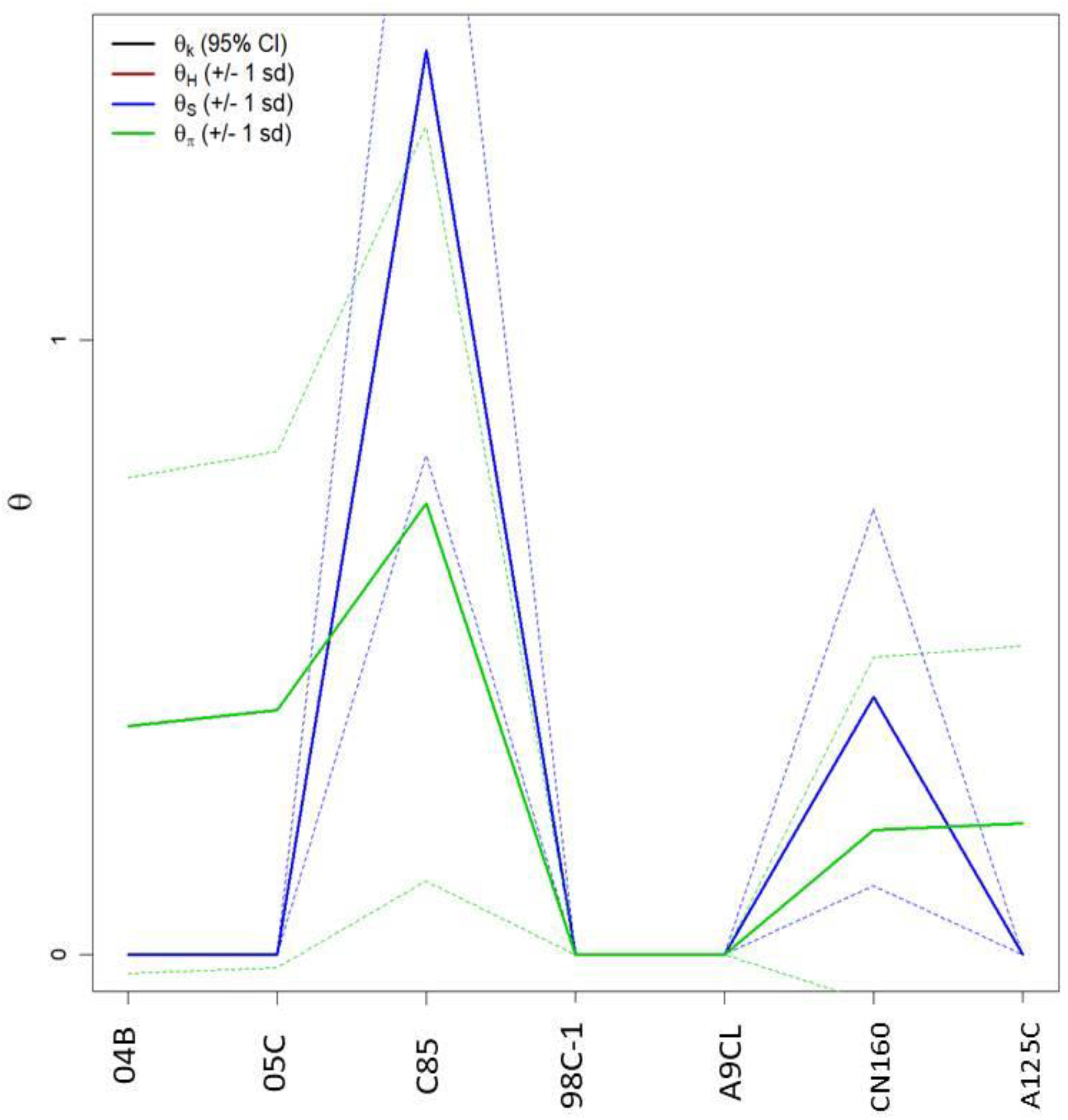
Graphic representation of molecular diversity indices in groups 04 B, 05 C, C 85, 98C-1, A9CL, CN160, A125C. *Generated by the statistical package in R language using the output data of the Arlequin software version 3.5.1.2.

**Figure 2b.**
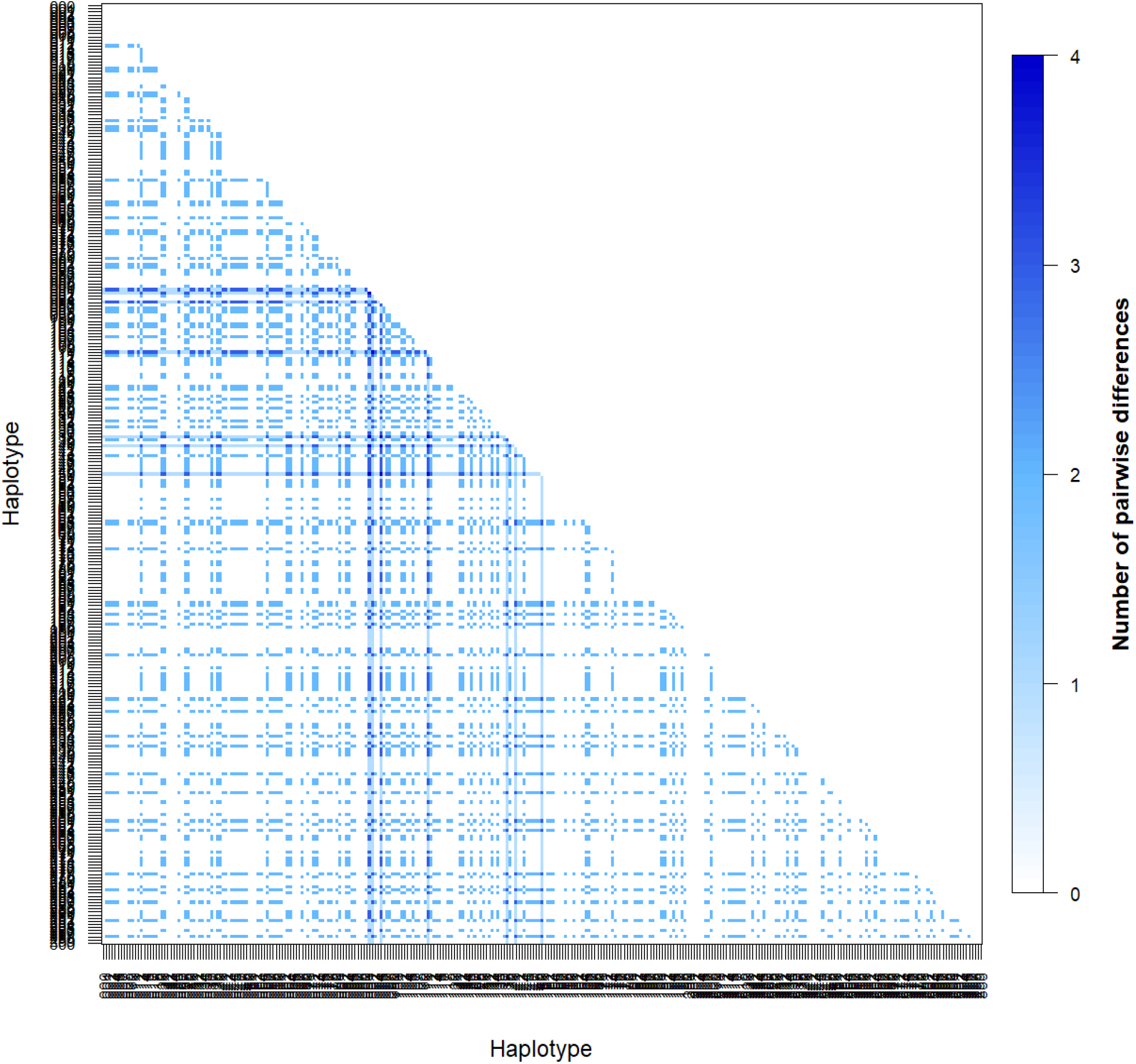
Representation of the haplotypic distance matrix among the 301 sequences studied. *Generated by the statistical package in R language using the output data of the Arlequin software version 3.5.1.2.

**Figure 3.**
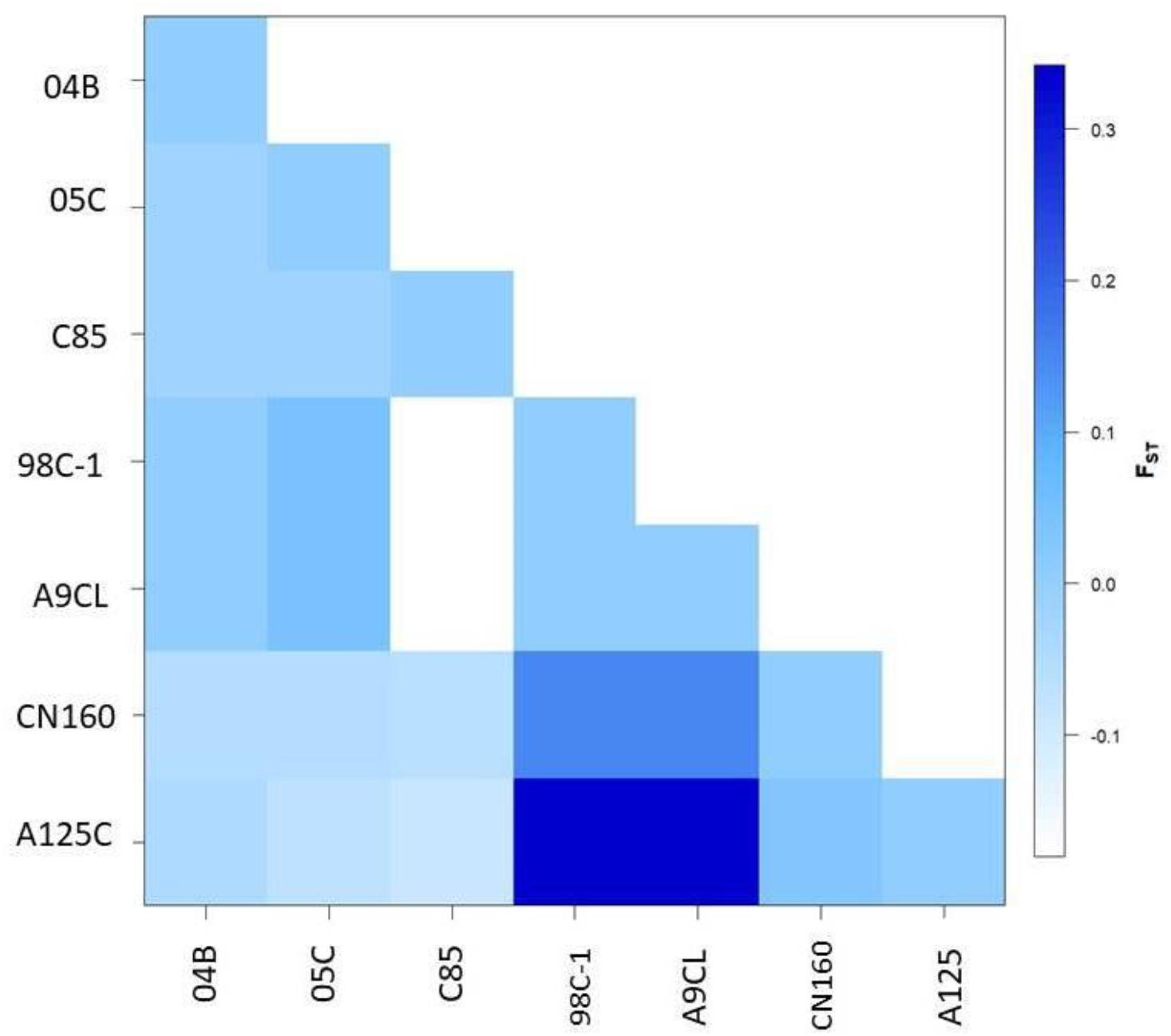
Matrix of genetic distance based on F_ST_ among the seven populations. * Generated by the statistical package in R language using the output data of the Arlequin software version 3.5.1.2.

**Figure 4.**
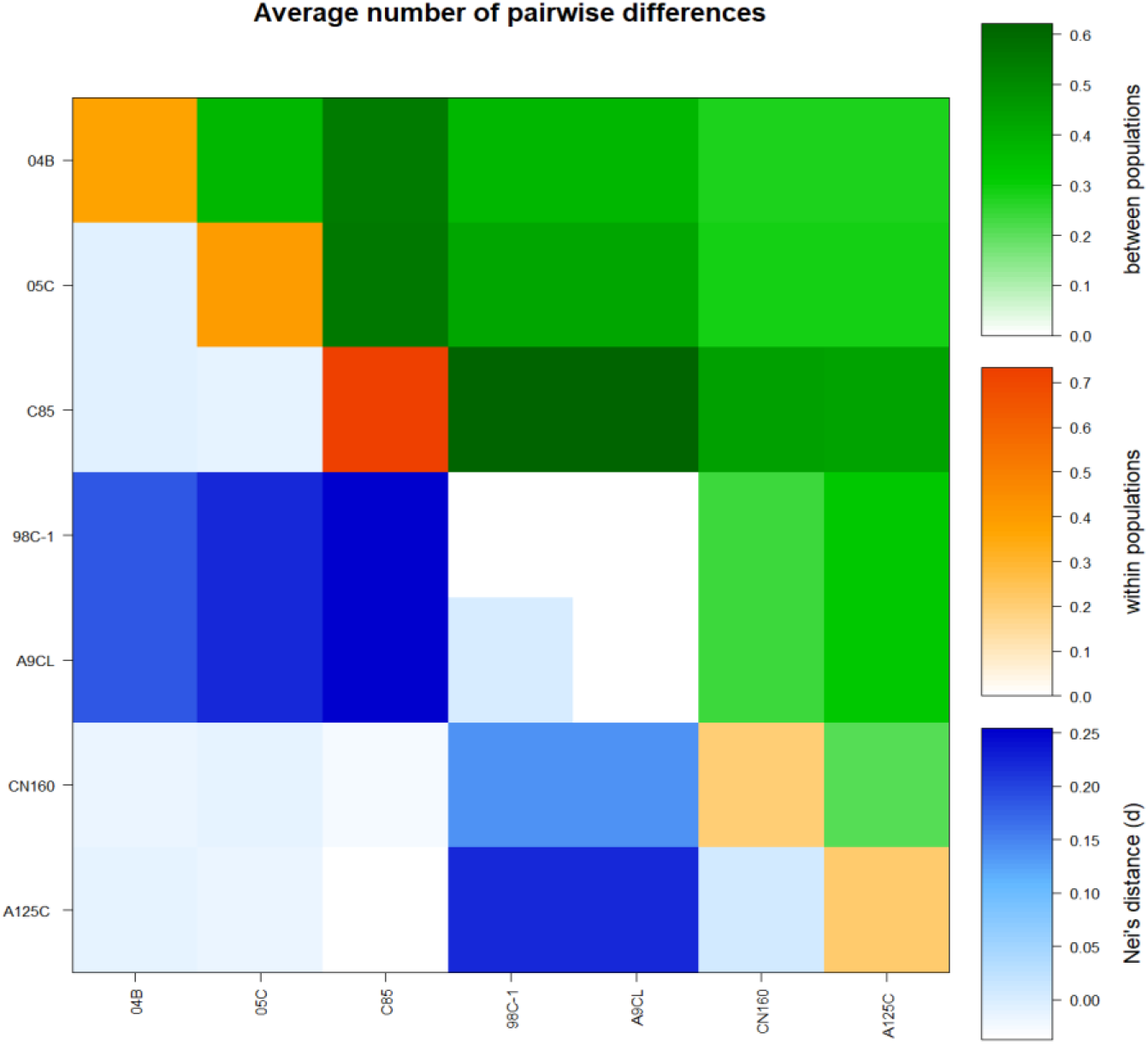
Matrix of paired differences between the populations studied: between the groups; within groups; and Nei distance for the seven groups. *Generated by the statistical package in R language using the output data of the Arlequin software version 3.5.1.2.

**Table 1.**
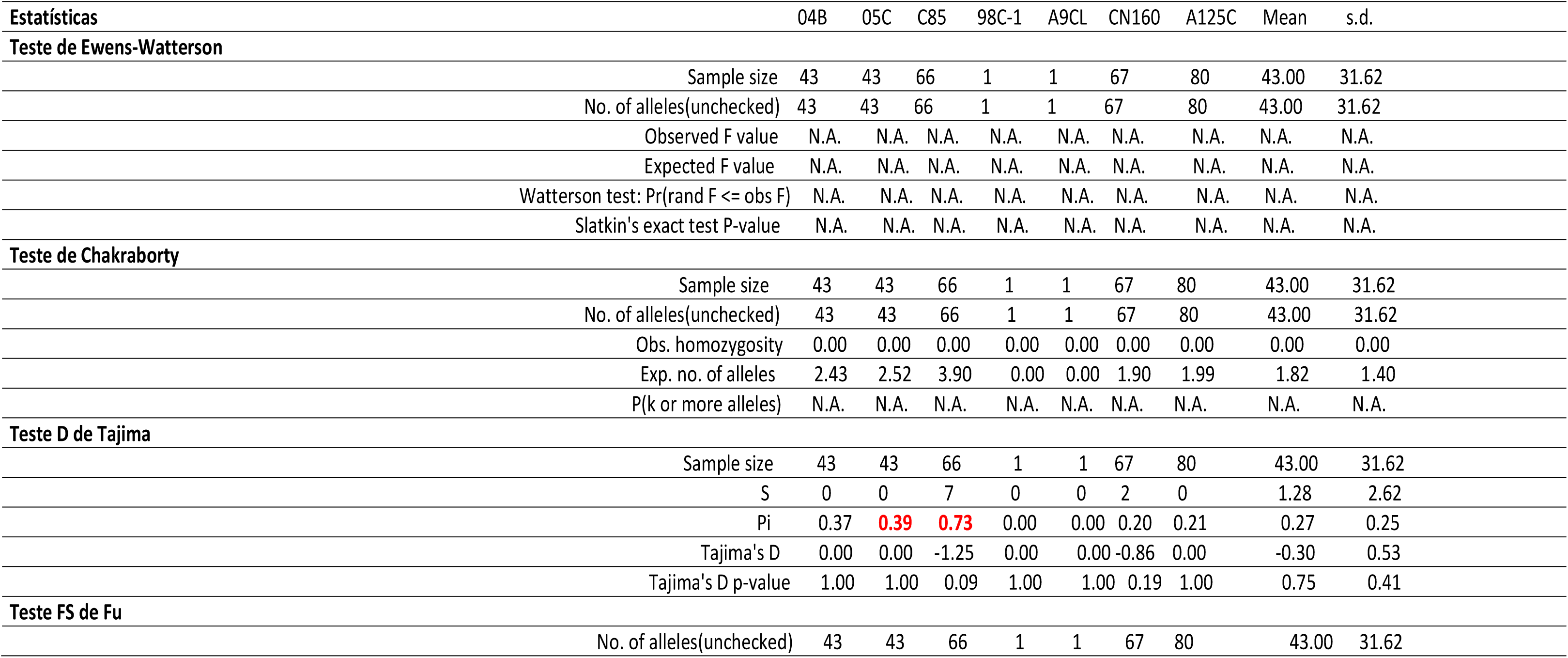
Neutrality test for the seven groups studied

The results presented suggest that the TP53 gene is a strong candidate in the construction of biosensors for the diagnosis of breast cancer in human populations, since its polymorphism levels are not significant and its molecular diversity indexes are unimpressive. Further analyses are still underway and we will soon have even more robust results corroborating our hypothesis.

## 7. Author’s contributions

To authors Xiao-Dan and collaborators by the availability of sequences in the public databank.

## 8. Acknowledgments

I thank my advisor Prof. Dr. Pierre Teodósio Félix; to UNIVISA for the availability and disposition of resources for the development of this work; to colleagues in the Lab for patience and to Laboratory of Population Genetics and Computational Evolutionary Biology - LaBECom.

